# RBPMS as a novel marker for cancer-associated fibroblasts in distinguishing left and right colorectal cancer sidedness

**DOI:** 10.1101/2025.08.05.668832

**Authors:** Sahira Syamimi Ahmad Zawawi, Ahmad Aizat Abdul Aziz, Andee Dzulkarnaen Zakaria, Lee Yeong Yeh, Nur Asyilla Che Jalil, Marahaini Musa

## Abstract

**Background:** Colorectal cancer (CRC) can be classified into left-sided (LCRC) and right-sided CRC (RCRC). These CRC entities present different molecular phenotypes and prognosis. Accumulation of cancer-associated fibroblasts (CAFs) reflects poor prognosis and recurrence of CRC. CAF-cancer cell crosstalk regulates fibroblast activation and drives CRC progression, facilitated via secretomes including TGFβ1. CAF-cancer cell interplay according to colon sidedness have yet to be fully investigated due to the lack of robust marker for CAF.

**Objective:** To explore a novel CAF marker in differentiating the mechanisms of LCRC and RCRC.

**Methods:** CAFs from cancerous tissues of LCRC and RCRC patients and normal fibroblasts from adjacent normal colon tissues were established. Profiling of these fibroblasts were performed. Gene expression of the fibroblasts was analyzed via Affymetrix Clariom S (Human) assay, validated using Western blot. The effect of TGFβ1 on protein expression of fibroblasts was also studied. Epithelial cancer cell lines were used as controls.

**Results:** Fibroblast profiling showed differences in morphology and proliferation between CAFs and NFs, and pro-proliferative effect of CAFs on CRC cells. Molecular analyses revealed RNA-binding protein with multiple splicing (*RBPMS*) to be differentially expressed between CAFs and NFs from left and right colon. *RBPMS* was significantly upregulated in CAFs from LCRC compared to their respective NFs. In contrast, *RBPMS* was downregulated in CAFs from RCRC compared to their NFs (*p*<0.05). Furthermore, TGFβ1 induced RBPMS expression in CAFs and NFs from the left colon, whereas it suppressed RBPMS expression in fibroblasts from right colon. CAFs from LCRC resembled myCAFs with myofibroblast-related expression signatures, whereas those from RCRC signified iCAFs with inflammatory marker expression.

**Conclusion:** This study highlighted RBPMS as a novel CAF marker in differentiating the mechanisms between LCRC and RCRC. This finding may be utilized in CAF-targeted therapy for CRC.

## Introduction

According to GLOBOCAN 2022, colorectal cancer (CRC) is the third leading cancer incidence affecting both males and females worldwide [1]. CRC is a complex disease. Two entities of CRC, namely left-sided CRC (LCRC) and right-sided CRC (RCRC) greatly differ based on their molecular phenotypes, prognosis, and treatment outcomes as summarized in Fig 1 [2]. This reflects the considerable heterogeneity of CRC by its sidedness, suggesting distinct underlying mechanisms.

**Fig 1.**
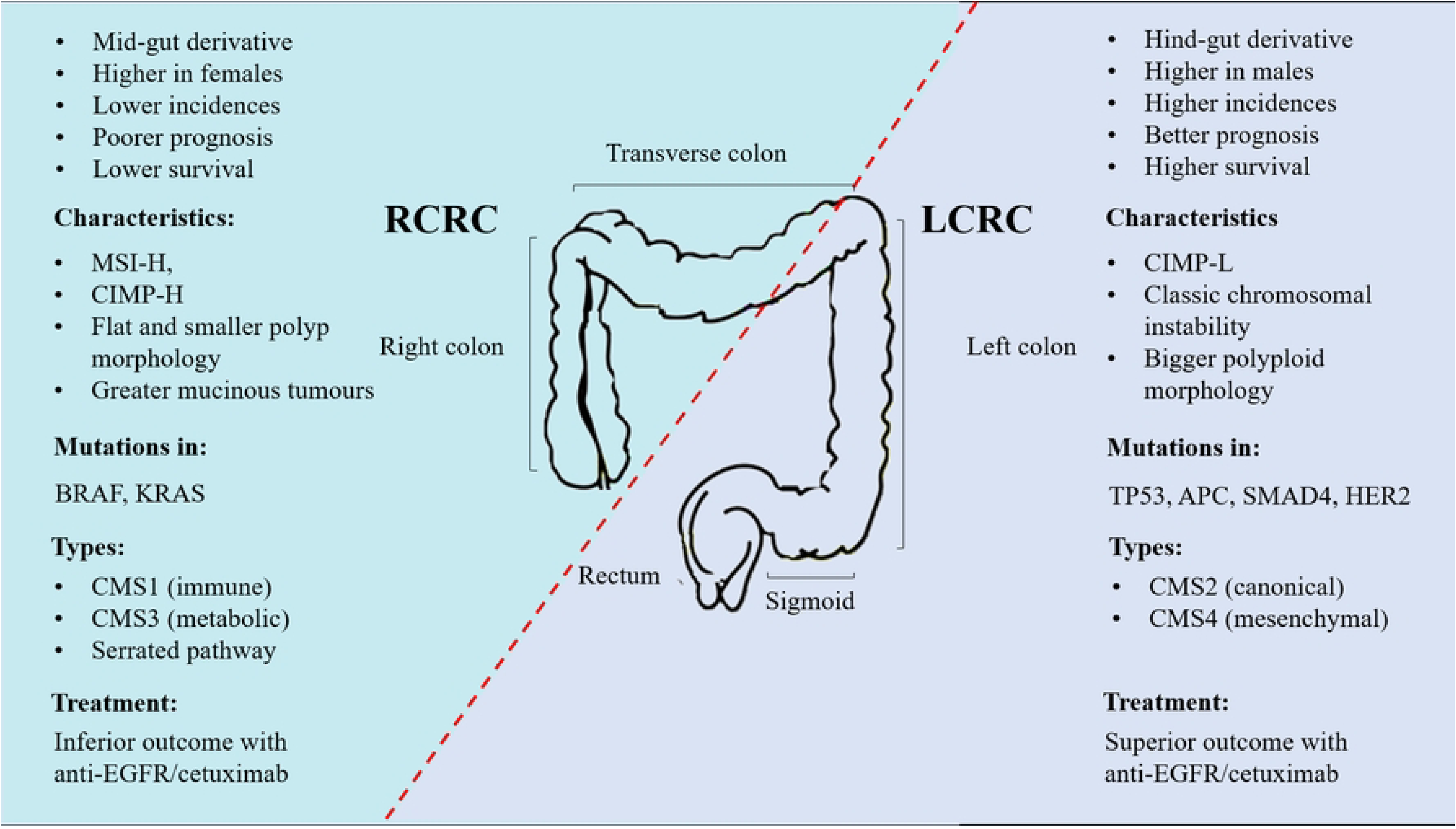
Differences between LCRC and RCRC.

Despite the advent of current therapy including the targeting immune checkpoint inhibitor, programmed cell death 1 (PD1) [3], 30-40% of CRC patients remain affected with increased 2-year recurrence [4]. This is partly attributed to cancer-associated fibroblasts (CAFs), the main cellular component in tumor microenvironment. Accumulation of CAFs indicates poor prognosis and recurrence of CRC [5]. CAF-cancer cells crosstalk facilitated by secretomes including transforming growth factor-beta 1 (TGF-β1) regulates fibroblast activation and drives colorectal carcinogenesis [6,7].

CAFs are multi-origin and may arise from trans-differentiation of non-activated fibroblasts [7]. CAFs can be recognized using conventional markers including alpha-smooth muscle actin (α-SMA), fibroblast-specific protein 1 (FSP1), and vimentin although their expressions are non-specific. Emerging CAF biomarkers including amine oxidase copper-containing 3 (AOC3) and leucine-rich repeat-containing 17 (LRRC17) were also reported [8, 9] although more studies warranted their performance as specific markers for CAF.

Many have investigated CAFs as a potential target for CRC treatment. Nonetheless, the influence of CAF-cancer cell crosstalk in regulating the mechanism of LCRC or RCRC has yet to be elucidated. Comprehensive investigation on CAFs with CRC sidedness is hindered by the lack of robust biomarkers. Hence, we aimed to explore a novel marker of CAFs that underlies the mechanistic differences between LCRC and RCRC. This will provide better insights toward tailoring future targeted approaches.

## Materials and method

### Established primary fibroblasts and epithelial CRC cell lines

Primary fibroblast cultures were established from 3 cm^2^ fresh resected tissues of LCRC (n=6) and RCRC (n=4) (denoted as CAFs, n=8), and their respective normal adjacent colon (normal fibroblasts, NFs, n=9). All samples were collected between 12 March 2022 and 16 March 2023 at Universiti Sains Malaysia (USM) Specialist Hospital, following written informed consent from all subjects and approval from Human Research Ethics Committee USM (code no: USM/JEPeM/20120685). The inclusion criteria were subjects aged ≥18, diagnosed with CRC, scheduled for first-time colorectal surgical resection, and treatment-naïve patients. Subjects with 2-years history of chemoradiotherapy for CRC and other cancer, resected part of the colon prior to surgery, pregnancy, and poor health status are excluded. Demographic and clinicopathological data of the subjects were collected.

For primary fibroblast establishment, CAFs and NFs were isolated from surgically resected colon tissues of CRC patients, based on gross and histological evaluation. Next, the acquired tissues were dissected and digested using a collagenase type IV (Thermo Scientific, USA) for 3 hours at 37°C, 5% CO2. The isolated cells were then grown in Dulbecco’s Modified Eagle Medium (DMEM) + 10% fetal bovine serum (FBS) + 3% penicillin-streptomycin solution. CAFs and NFs of passage numbers of 3 to 10 were used in subsequent experiments to avoid replicative senescence.

Two epithelial cell lines derived from CRC and cervical cancer namely SW620 and C33A, respectively were obtained from American Type Culture Collection (ATCC) and included as controls. Cells from different origins were selected to compare the target protein expression in different organs. These epithelial cell lines were grown in DMEM +10% FBS + 1% penicillin-streptomycin. Cells were maintained in a humidified incubator (37°C, 5% CO2).

Cell morphology was observed under an inverted light microscope and captured by using ToupView Camera software. Cells were checked for mycoplasma contamination. Cell lines were passaged regularly once reached 80-90% confluency and stored in a liquid nitrogen tank for long term storage.

For fibroblast profiling, the primary fibroblast morphology was observed and compared between CAFs and NFs derived from left and right colon. In addition, MTT assay was performed to study proliferative activity of the CAF and NF lines, as described in the next section.

### 3-(4,5-dimethylthiazol-2-yl)-2,5-diphenyl-2H-tetrazolium bromide (MTT) proliferation assay

MTT assay was conducted to assess a) the proliferation of CAFs and NFs in culture and b) their influence on SW620 proliferation. The following terms are used to refer to specific groups of the fibroblasts; ‘R-CAF’ refers to CAFs from RCRC, ‘L-CAF’ to CAFs from LCRC, ‘R-NF’ to NFs from right-sided normal, and ‘L-NF’ to NFs from left-sided normal.

For fibroblast growth profiling, CAF and NF lines were seeded at 1×10^3^ and incubated for 24, 48, 72, and 96 hours with a complete culture medium (DMEM + 10% FBS + 3% penicillin-streptomycin solution). Then, the cells were treated with MTT reagent (ab211091, Abcam, USA) for 3 hours and 15 minutes. The optical density (OD) was obtained from absorbance value at λ590nm using a spectrophotometer (Varioskan Flash, Thermo Scientific, USA). CAFs and NFs proliferation were later determined and compared.

To study the effect of the fibroblasts on SW620 proliferation, SW620 cells were seeded in a 96-well plate at 5×10^3^ and 10×10^3^, incubated at 37°C, 5% CO_2_ for 24 hours and subsequently treated with conditioned medium (CM) of either CAFs (CM-L-CAF or CM-R-CAF) or NFs (CM-L-NF or CM-R-NF) for 72 hours. CM of fibroblasts was prepared by incubating CAFs and NFs at 85% confluency with DMEM alone (serum-free (SF) medium) for 48 hours at 37°C, 5% CO_2_ before the medium was collected and used in the experiment. A similar MTT assay protocol as described previously was performed. SW620 proliferation under CM treatment from CAFs and NFs was determined using the following formula:

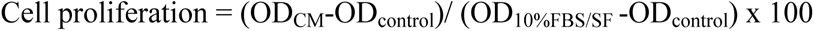

D_CM_ = Absorbance of cells treated with CM of CAFs or NFs from different colon sidedness
D_10%FBS/SF_ = Absorbance of cells treated with DMEM containing 10% FBS or SF medium
D_control_ = Absorbance of control cells, treated with SF medium

### Microarray analysis

A total of 2×10^6^ of CAFs and NFs from LCRC and RCRC, and 1×10^6^ of SW620, were seeded in 75cm^2^ and 25cm^2^ flasks respectively. Total RNA was isolated from these cells using a RNeasy® Mini Kit (cat. 74106, Qiagen, USA) according to the manufacturer’s instructions with several modifications. RNA samples (n=8) with satisfactory concentration and purity tested with Nanodrop spectrophotometry (Multiskan^TM^ SkyHigh, Thermo Scientific, USA) were selected for further analysis. Gene expression analysis was performed via Affymetrix Clariom S (Human) assay (Thermo Fisher) (Research Instruments, Malaysia). Arrays were stained using the FS450_0007 fluidics protocol and screened using a GeneChip™ Scanner 3000 7G (Applied Biosystems, Thermo Scientific, USA). The data was then imported into Transcriptome Analysis Console operating on R/Bioconductor [10]. Data was normalized via robust multiarray average algorithm background correction. Differentially expressed genes (DEGs) were identified within the threshold fold change>2 or <-2 and *p*-value < 0.05, adjusted using Benjamini-Hochberg procedure. The *p*-value was transformed to -log10 scale for visualization in volcano plots. The data reported in this article have been deposited in the Gene Expression Omnibus database, www.ncbi.nlm.nih.gov/geo (accession no. GSE274061).

### Western blot analysis

Cells were seeded at the same densities as described in the **Microarray analysis** section. Cells were lysed using M-PER lysis buffer (Thermo Scientific, USA) and centrifuged at 12000 rpm, 4°C. Protein lysates were collected. Protein concentration of the lysate was assessed using Pierce BCA assay (cat. 23225, Thermo Scientific, USA).

Proteins were Western blotted and later the blots were blocked with 1% Bovine Serum Albumin in PBS for 40 minutes at room temperature (RT). Then, the membranes were incubated overnight at 4°C with selected primary antibodies in the blocking solution. The blots were then incubated with secondary antibodies for 2 hours at RT and washed with 1X TBS-T. Immunoreactive bands were visualized using Fluor ChemM Western Blot Imager (ProteinSimple, USA) after incubation with Clarity^TM^ Western ECL (cat. 170-5061, Bio-Rad, USA). Band intensities were analyzed with Image J1.53k software.

Selected primary antibodies were as follows: RBPMS (1:500, ab152101, Abcam, USA), α-SMA (1:10000, ab7817, Abcam, USA), AOC3/VAP-1 (1:200, sc-33670, Santa Cruz Biotechnology, Europe), LRRC17 (1:500, VK3114774A, Invitrogen, Sweden), EpCAM (1:500, ab223582, Abcam, USA), and antibody β-tubulin (1:150, sc-555329, Santa Cruz, Europe). The horseradish peroxidase-conjugated secondary antibodies included were donkey anti-rabbit (1:2000, ab16284, Abcam, USA) and goat anti-mouse (1:5000, ab205719, Abcam, USA).

### TGFβ1 treatment

To study the effect of TGFβ1 on CAFs and NFs, a total of 2×10^6^ fibroblasts were seeded and treated with 10ng/ml TGFβ1 (Peprotech) prepared in SF medium (i.e. DMEM + TGFβ1) for 72 hours. The proteins expression in the fibroblasts was analyzed via Western blot analysis. Morphology of the TGFβ1-treated fibroblasts was also recorded.

### Statistical analysis

All data(s) except from microarray analysis are presented as mean ± SEM from duplicate experiments. GraphPad Prism were employed for all statistical analyses. Paired two-tailed t-test was used for comparison between the two respective groups. The *p*-value<0.05 shows statistical significance and *p*<0.01 indicates a higher statistical significance.

## Results

### Different phenotypic characterization of primary fibroblasts according to colon sidedness

CRC patients’ clinicopathological characteristics are included in Table 1. L-CAF and R-CAF were found to be larger in size and have nuclei with thicker elongated bodies compared to their respective NFs, which resembled more spindle-shaped morphology (Fig 2A-B). Based on MTT assay, L-CAF and R-CAF demonstrated greater proliferation than their respective NFs (Fig 2C-D). Interestingly, R-CAF showed significantly higher proliferation than L-CAF (*p*<0.05).

**Fig 2.**
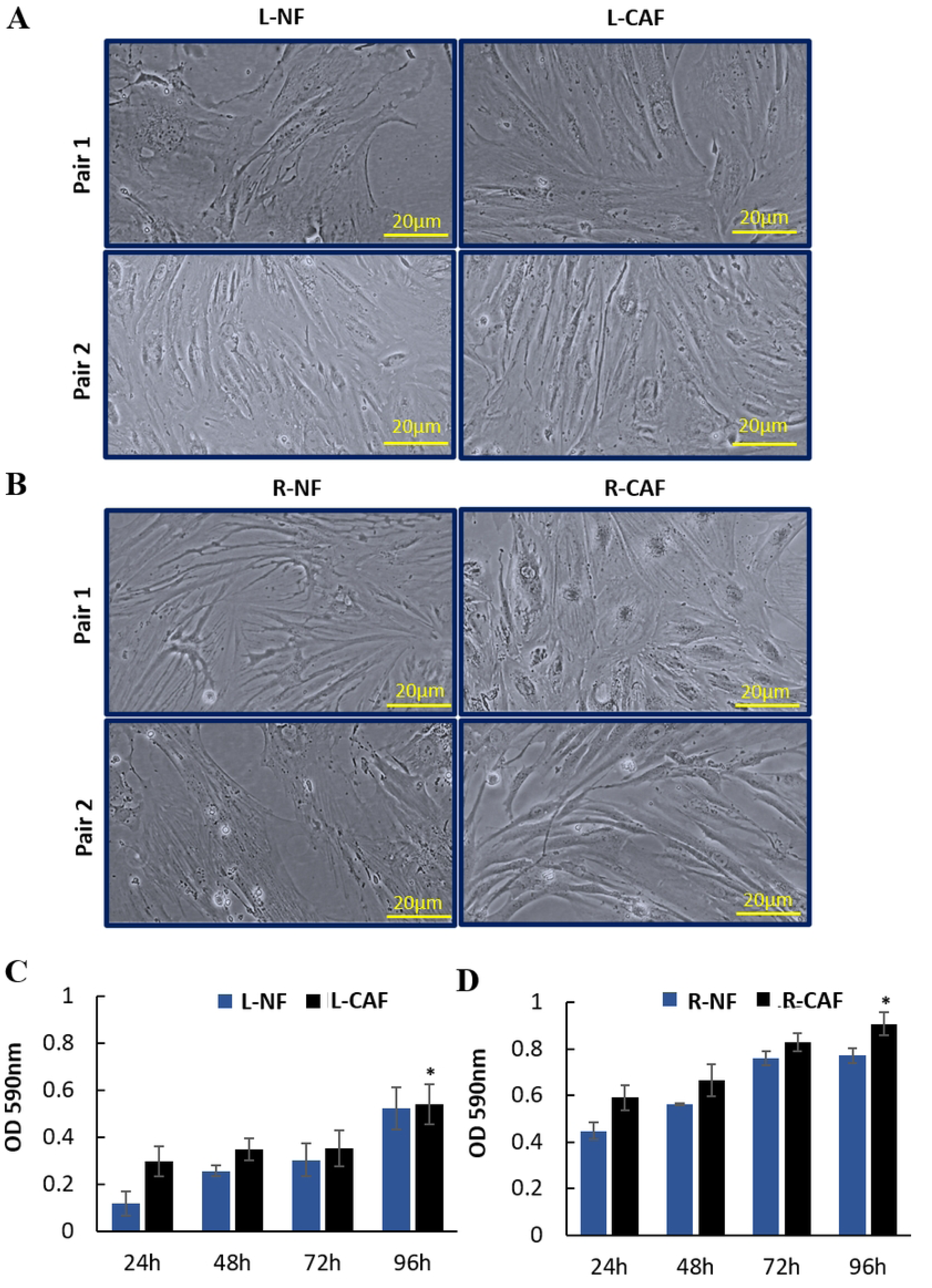
Characterization of primary fibroblasts. (A-B) Cells morphology. (C-D) Proliferation activity of cells after incubation for different time points. Data(s) are presented as mean (absorbance (OD) ±SEM). \**p*<0.05 compared between L-CAF and R-CAF, two-tailed paired t-test.

**Table 1.**
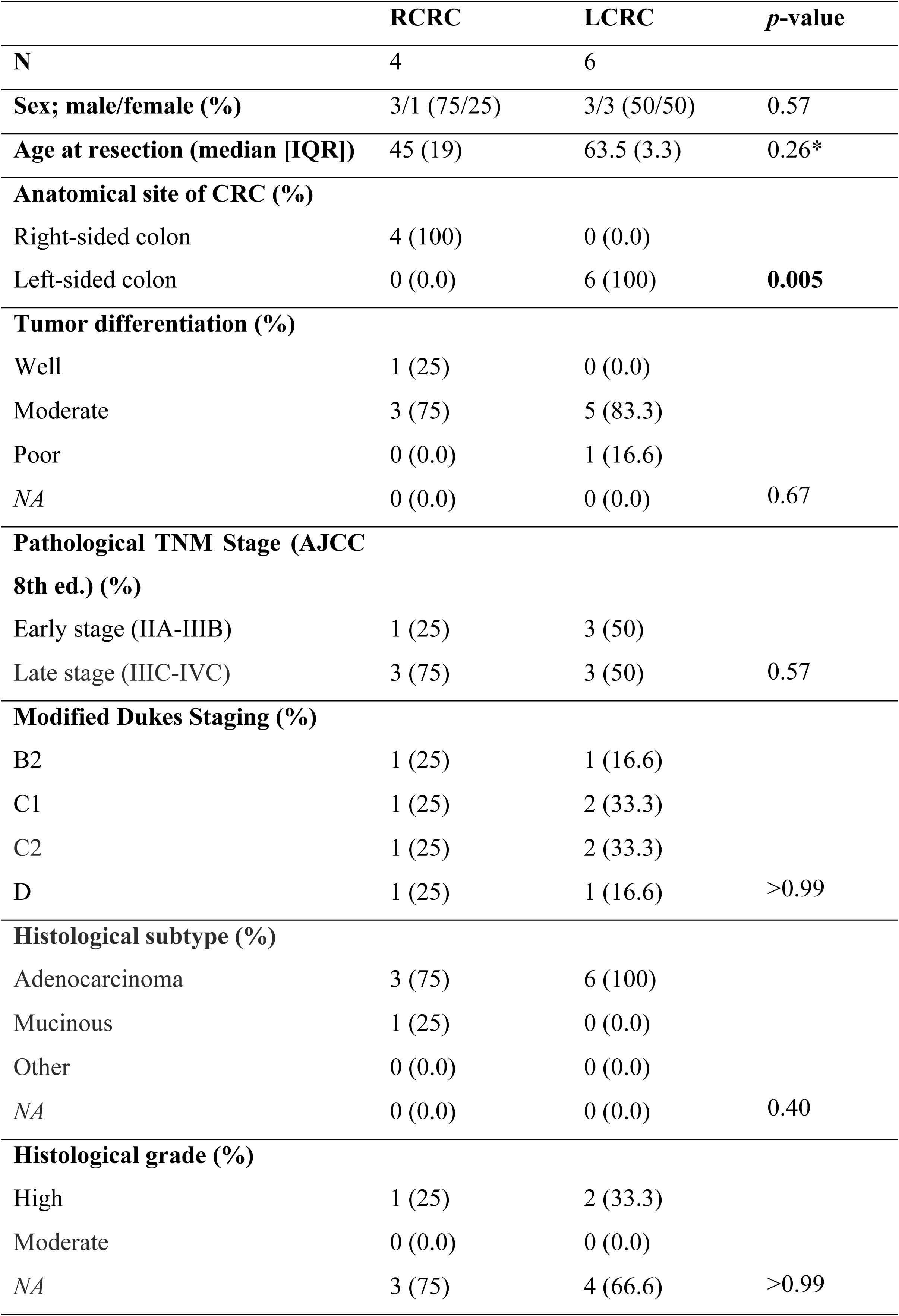
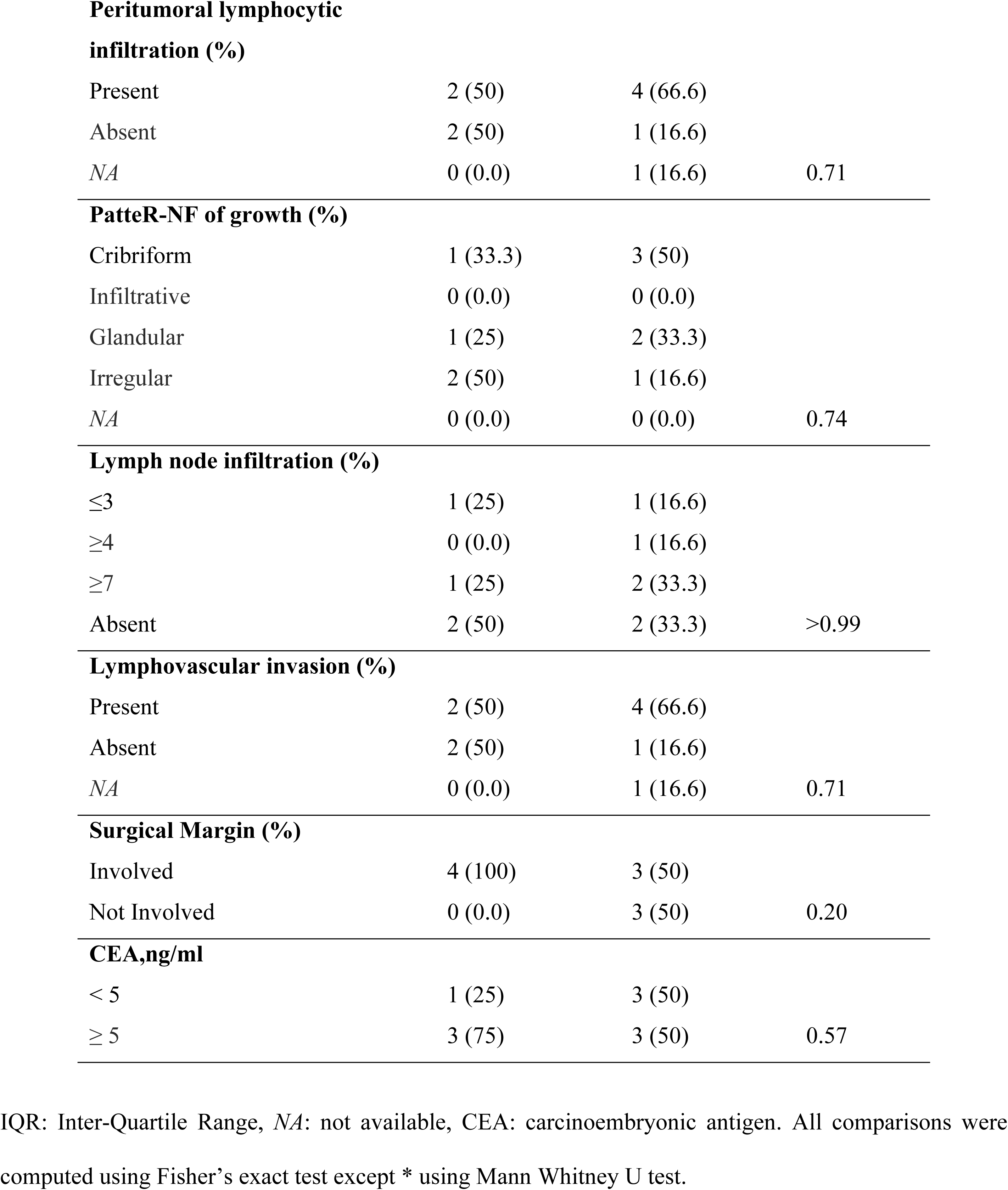
CRC patients clinicopathological characteristics.

### CAF-SW620 cells crosstalk promote significant SW620 proliferation

CM-L-CAF and CM-R-CAF induced greater proliferation of SW620, with a significant difference to SF medium (*p*<0.01) (Fig 3A-B). Conversely, SW620 cells showed a significant decline of proliferation when incubated with CM-L-NF and CM-R-NF.

**Fig 3.**
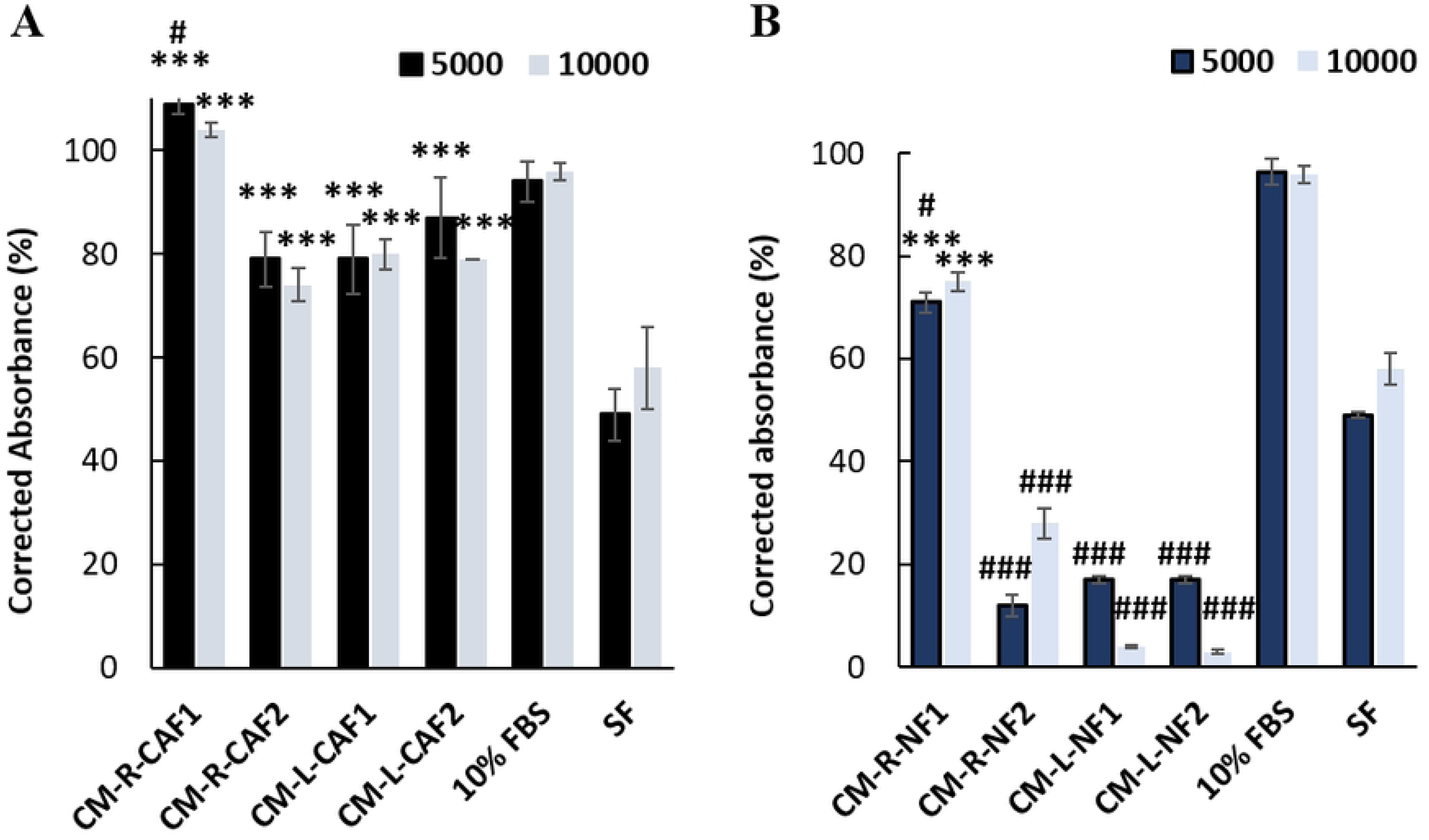
Proliferation of SW620 cells upon treatment with (A) CM-R-CAF and CM-L-CAF, and (B) CM-R-NF and CM-L-NF. Data(s) are presented as mean (corrected absorbance (%) ± SEM). \*\*\**p*<0.01 compared with cells treated with SF medium (negative control); #*p*<0.05, ###*p*<0.01 compared with cells treated with DMEM supplemented with 10% FBS (positive control), two-tailed paired t-test.

### Different characterization of primary fibroblasts relative to SW620

Microarray analysis of primary fibroblasts vs SW620 revealed 1595 DEGs; 1398 (upregulated) and 197 (downregulated). Caldesmon1 (*CALD1*), decorin (*DCN*), secreted protein acidic and cysteine-rich (*SPARC*), collagen type (Iα1-chain) (*COL1A1*), (XIIα1-chain) (*COL12A1*), and RNA-binding protein with multiple splicing (*RBPMS*) were significantly upregulated in fibroblasts comparative to SW620 (*p*<0.05) (Fig 4A-B) (Table 2). Conversely, epithelial markers including keratin 18 (*KRT18*) and epithelial cellular adhesion molecule (*EpCAM*) were significantly downregulated in fibroblasts in comparison to SW620.

**Fig 4.**
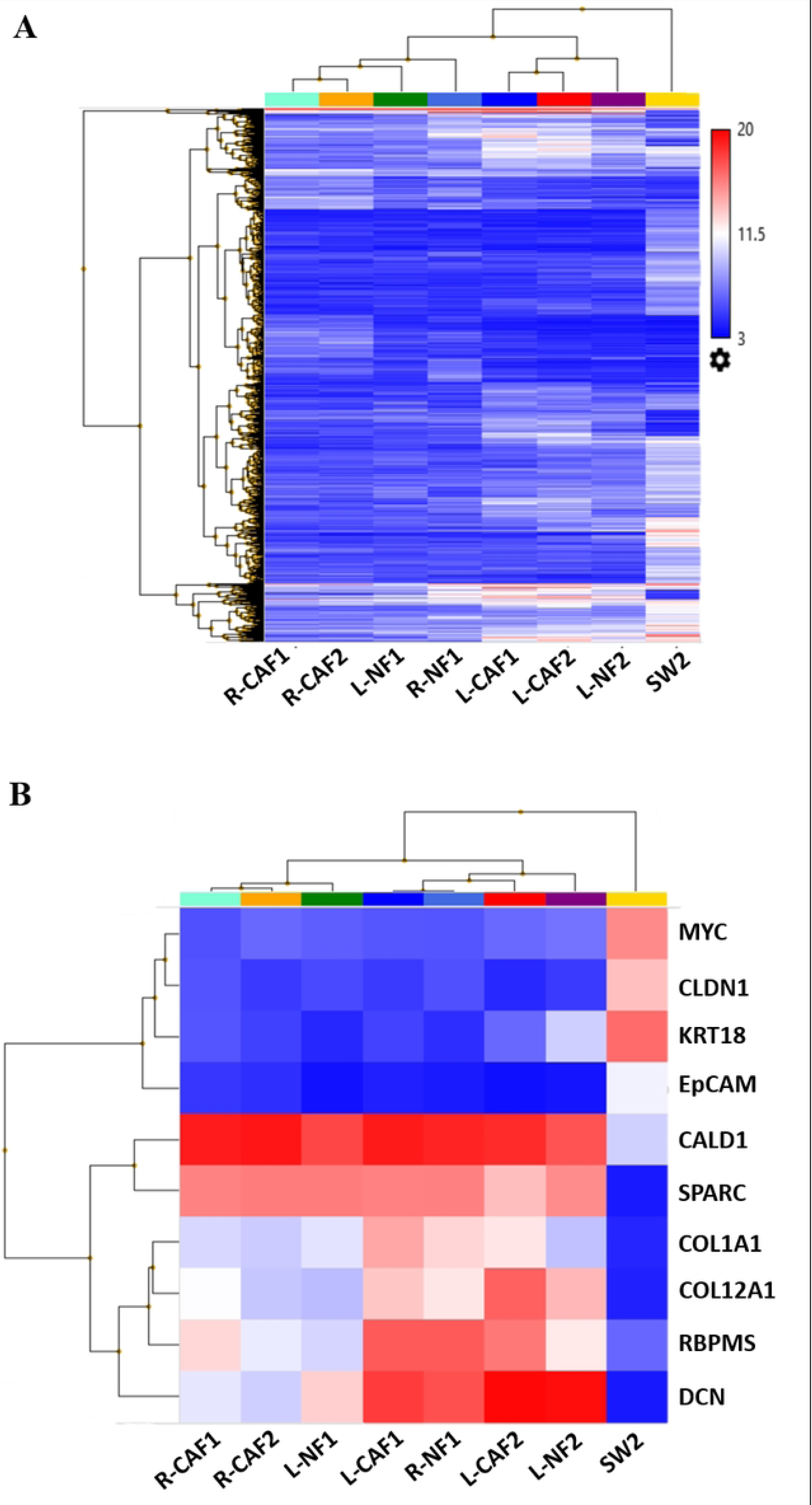
Microarray analysis between fibroblasts and SW620 (denoted as SW2). (A) Overall heatmap representation. (B) Top 10 DEGs between fibroblasts and SW620. Blue denotes gene downregulation; red denotes gene upregulation.

**Table 2.**
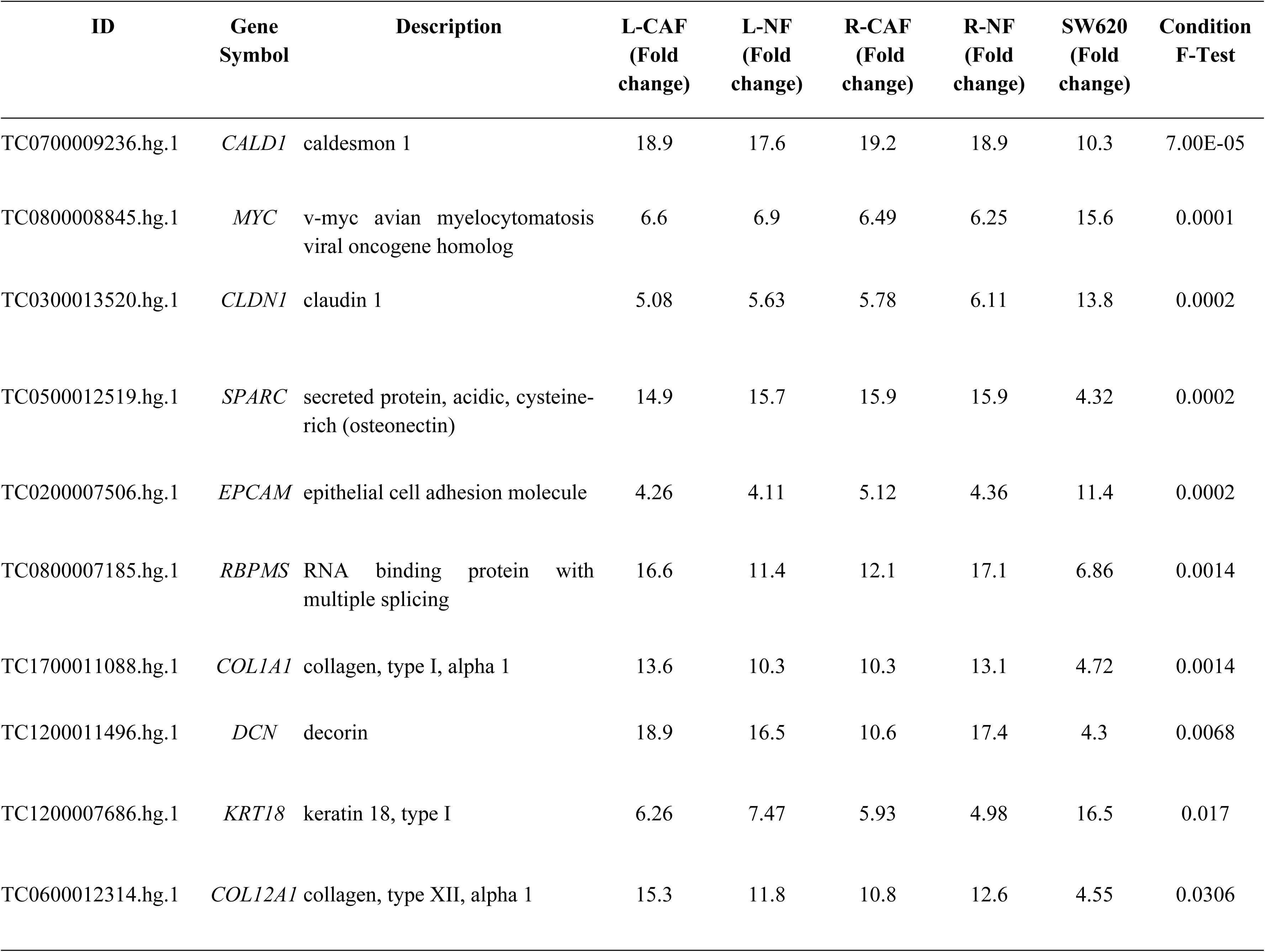
Top 10 DEGs between fibroblasts and SW620.

### Different CAF signatures between RCRC and LCRC

Microarray analysis of L-CAF vs R-CAF was computed which revealed 2035 DEGs; 960 (upregulated), 1075 (downregulated). L-CAF exhibited significant upregulation of myofibroblast-related markers; *COL1A2*, actinin alpha 1 (*ACTN1)*, actin alpha 2, smooth muscle (*ACTA2)* the encoding gene for α-SMA, *TGFβ1* and ECM-myofibroblast markers; thrombospondin 1 (*THBS1)*, matrix metalloproteinase 2 (*MMP2),* and insulin-like growth factor binding protein 5 (*IGFBP5*) (Fig 5A). Conversely, R-CAF was enriched with inflammatory markers including *IL6* and several CCL and CXCL motif cytokines. Microarray analysis of L-NF vs R-NF was also computed which revealed 562 DEGs; 430 (upregulated) 132 (downregulated) (Fig 5B).

**Fig 5.**
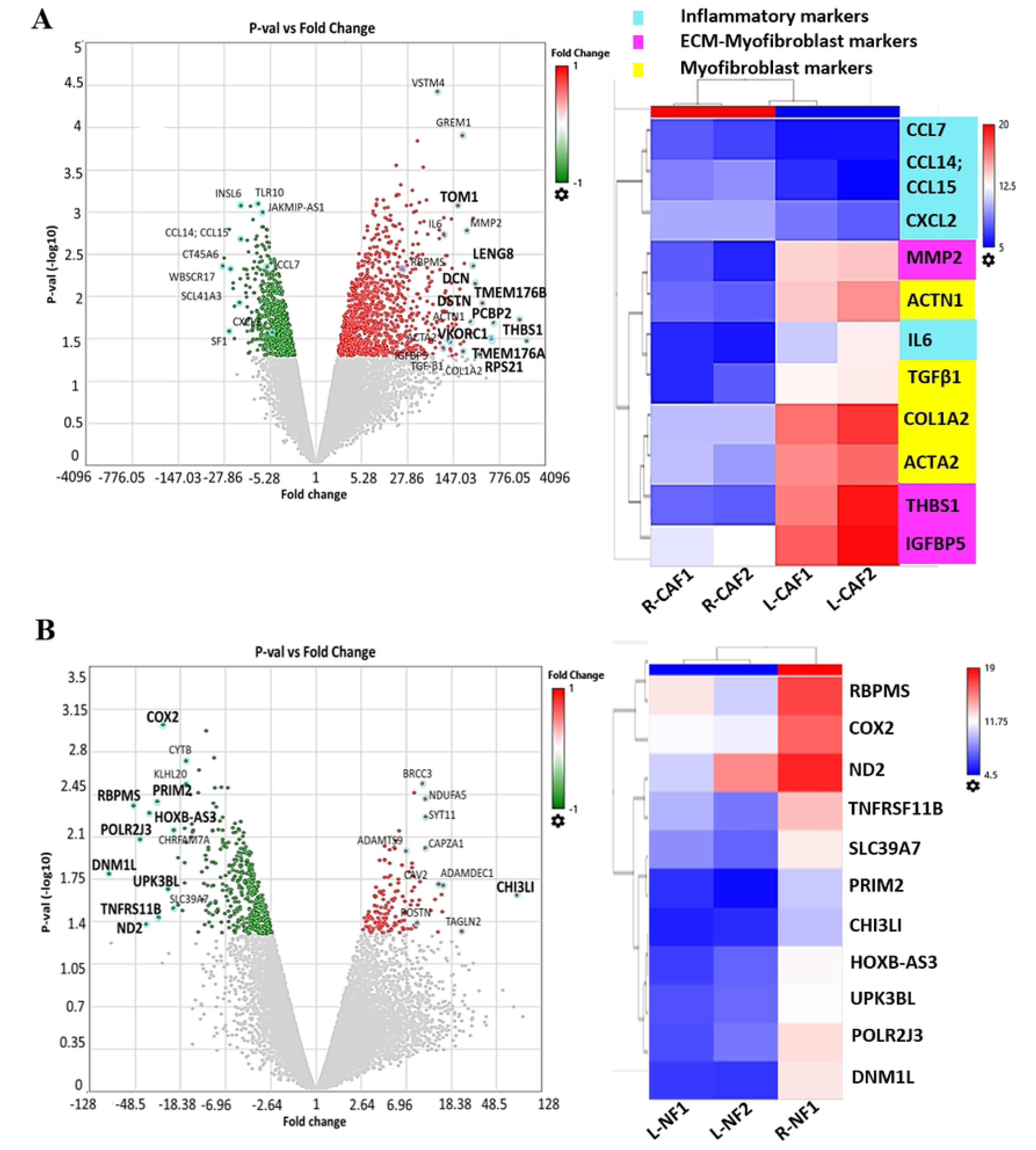
Microarray analyses of (A) L-CAF vs R-CAF, represented by volcano plot and heatmap of different markers expression, and (B) L-NF vs R-NF, represented by volcano plot and heatmap of top 10 DEGs. In the volcano plots, red, green, and gray points indicate upregulated genes, downregulated genes, and non-differentially expressed genes, respectively. Significant DEGs are shown in blue circles and the top 10 DEGs are indicated in bold. In heatmaps, blue denotes gene downregulation; red denotes gene upregulation.

### RBPMS as a novel CAF marker in delineating mechanisms between RCRC and LCRC

*RBPMS* was identified for its high expression in fibroblasts compared to SW620 and as one of the top shared DEGs between different fibroblast groups (Fig 6A-B). There was a stark contrast of *RBPMS* expression across different fibroblast groups (Fig 7). Interestingly, *RBPMS* was revealed to be upregulated in L-CAF compared to L-NF, whereas it was downregulated in R-CAF compared to R-NF. Furthermore, *RBPMS* was shown to be downregulated in L-NF in comparison to R-NF, whereas it was upregulated in L-CAF in comparison to R-CAF.

**Fig 6.**
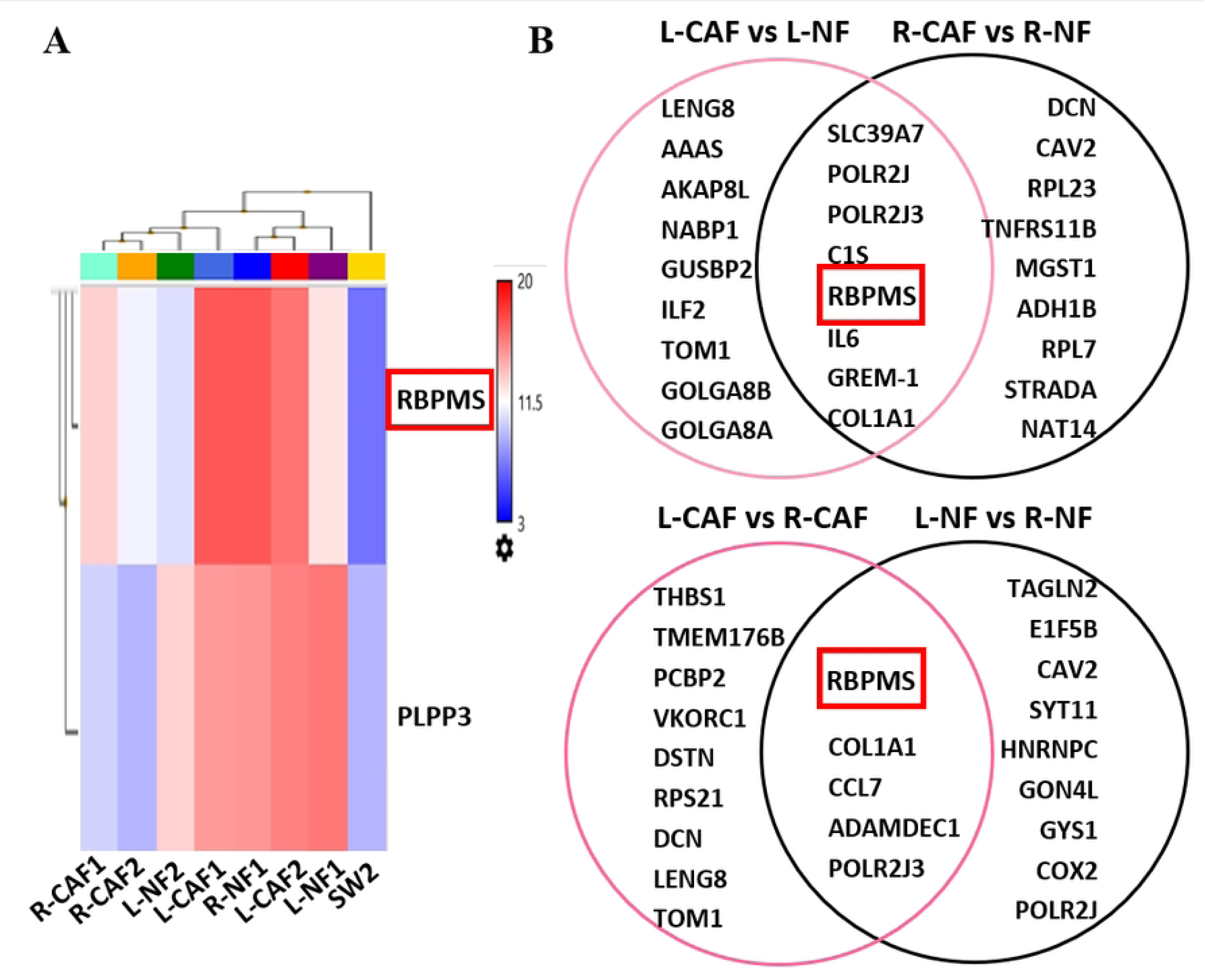
RBPMS specific expression in differentiating (A) fibroblasts and SW620, represented by heatmap (blue denotes gene downregulation, red denotes gene upregulation), and (B) fibroblasts from different colon sidedness, represented by Venn diagrams.

**Fig 7.**
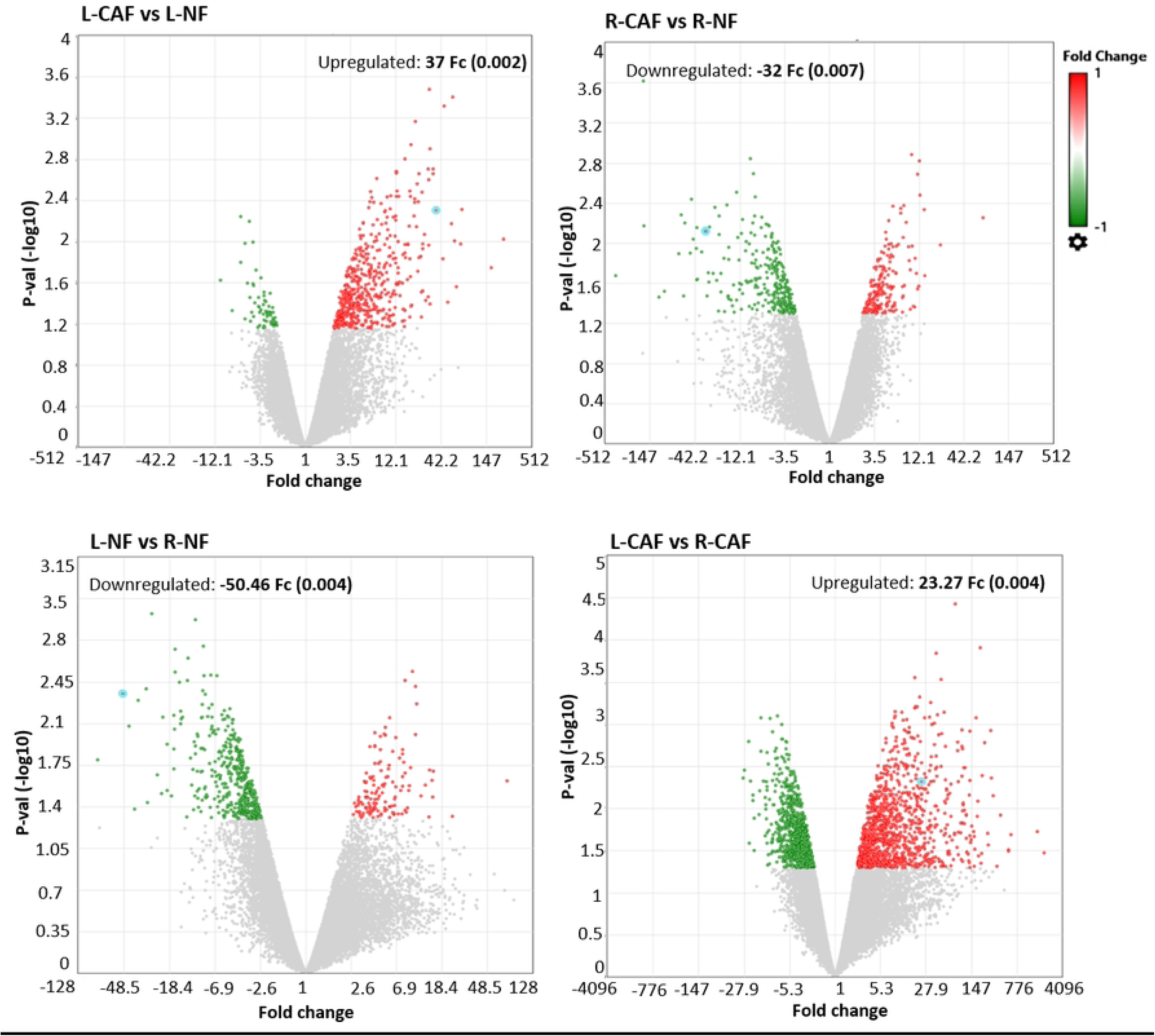
Volcano plots illustrating DEGs between different fibroblast groups. RBPMS differential expression is demonstrated in blue circle. Fc: Fold change.

Further analysis confirmed the RBPMS positive expression in selected fibroblast lines (Fig 8A). Faint bands were observed for SW620, which coincide with the information from the Protein Atlas database with normalized transcript expression per million (nTPM) 16.6 (http://www.proteinatlas.org/). No band was observed in C33A, consistent with nTPM data. The specific RBPMS expression between CRC entities derived from microarray analysis was validated at the protein level with statistical significance (*p*<0.05, Fig 8B).

**Fig 8.**
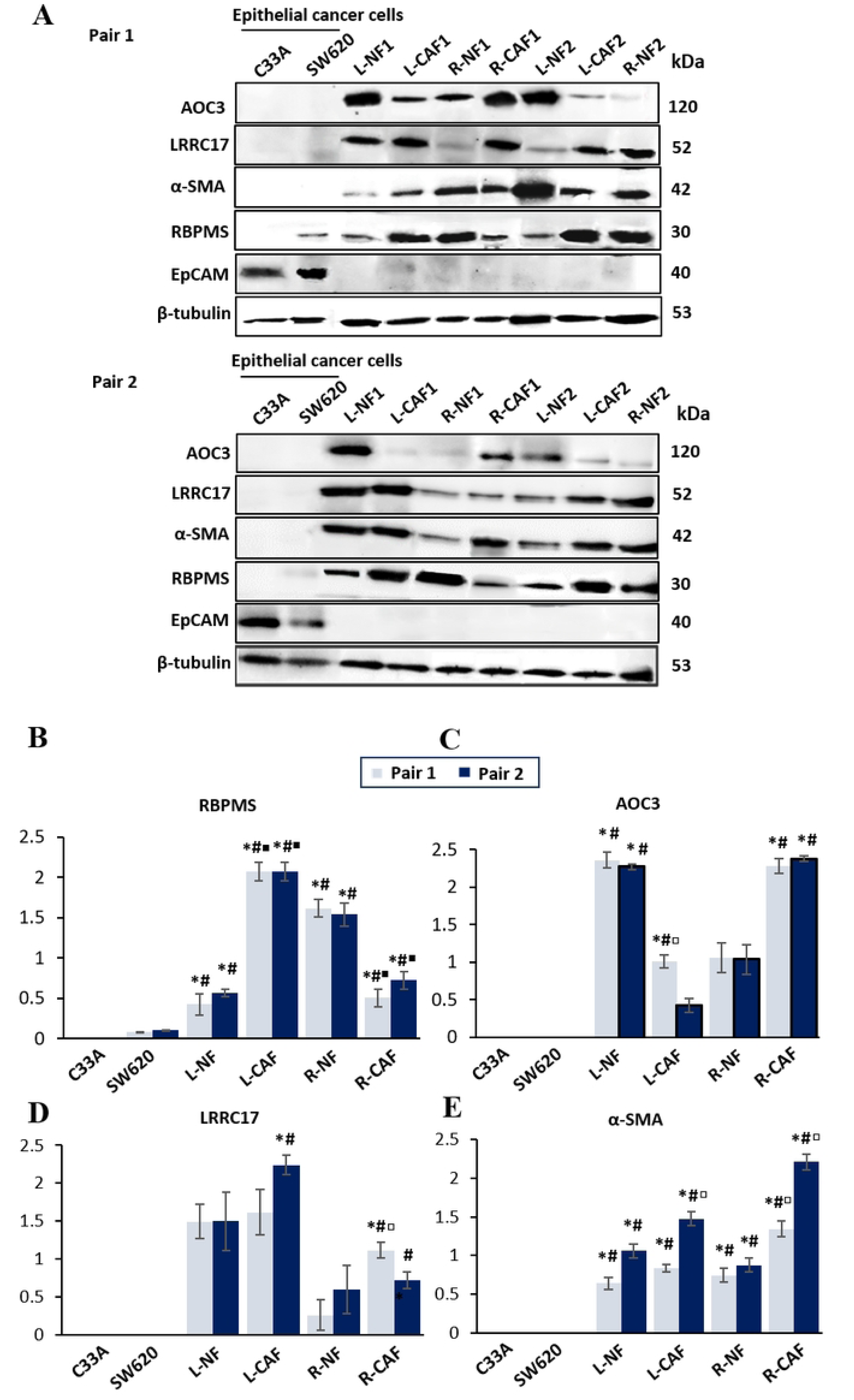
Western blot analyses between fibroblasts and epithelial cancer cells. (A) Images of bands. (B) Expression of RBPMS, (C) AOC3, (D) LRRC17, and (E) α–SMA, relative to β-tubulin. Data are presented as mean ± SEM from duplicate experiments. \**p*<0.05 compared with C33A (control), #*p*<0.05 compared with SW620 (control), ▫*p*<0.05 compared with respective NFs only, ▪*p*<0.05 compared between L-CAF and R-CAF, and with respective NFs.

### AOC3, LRRC17, and α-SMA expression characterize activated fibroblasts

Further characterization of fibroblasts was done by detecting α-SMA, AOC3, and LRRC17 expression via Western blot. AOC3, LRRC17, and α-SMA were positively expressed in all fibroblasts except epithelial cells (Fig 8A). Contrarywise, EpCAM, an epithelial marker, was positively expressed in epithelial cancer cells but negatively expressed in the fibroblasts. This validated the microarray result of *EpCAM* upregulated expression in SW620 comparative to fibroblasts.

For AOC3, the significant relative expression only showed in L-NF vs L-CAF of Pair 2 (Fig 8C). For LRRC17, only R-CAF of both pairs and L-CAF of Pair 2 showed significant difference when compared to epithelial cells (Fig 8D). α-SMA expression, indicative of activated fibroblasts, was higher in fibroblasts than the epithelial cells (Fig 8E).

### TGFβ1 signaling influence the specific expression of RBPMS in CRC progression

RBPMS was regulated differently in the TGFβ1-treated fibroblasts, based on colon sidedness (Fig 9A). TGFβ1 significantly induced RBPMS expression in L-CAF and L-NF, whereas suppressed RBPMS in R-CAF and R-NF (*p*<0.05) (Fig 9B). AOC3 downregulation was detected in the TGFβ1-treated L-NF and R-CAF when compared to DMEM+10% FBS groups (Fig 9C). Contrarywise, α-SMA and LRRC17 upregulation were observed in TGFβ1-treated fibroblasts regardless of the colon sidedness (Fig 9D-E). A slight difference in morphology of TGFβ1-L-NF (Fig 10) was observed where these cells display thicker elongated cell bodies and more prominent indented nuclei, mimicking the myofibroblasts, different to those L-NF grown in DMEM+10% FBS.

**Fig 9.**
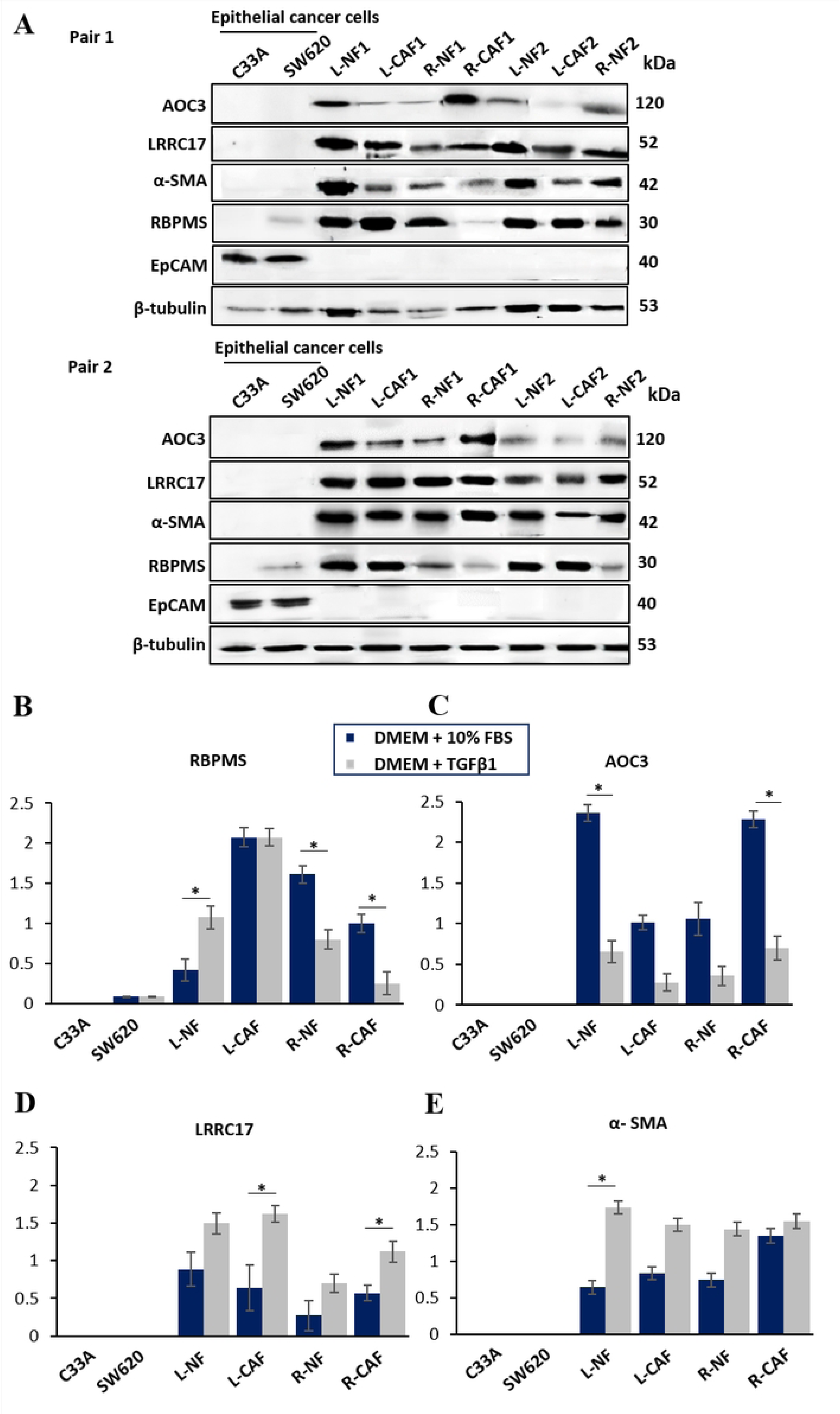
Western blot analysis of fibroblasts following 72 hours treatment under DMEM + TGFβ1 (10 ng/mL). (A) Images of bands showing protein expression following TGFβ1 treatment. (B) Expression of RBPMS, (C) AOC3, (D) LRRC17, and (E) α-SMA, relative to β-tubulin. Data are presented as mean ± SEM from two independent experiments performed in duplicate, comparing fibroblast groups treated with DMEM + 10% FBS and DMEM + TGFβ1. \**p*<0.05 compared with C33A (control), #*p*<0.05 compared with SW620 (control), ▫*p*<0.05 compared with respective NFs, ▪*p*<0.05 compared between L-CAF and R-CAF, and with respective NFs.

**Fig 10.**
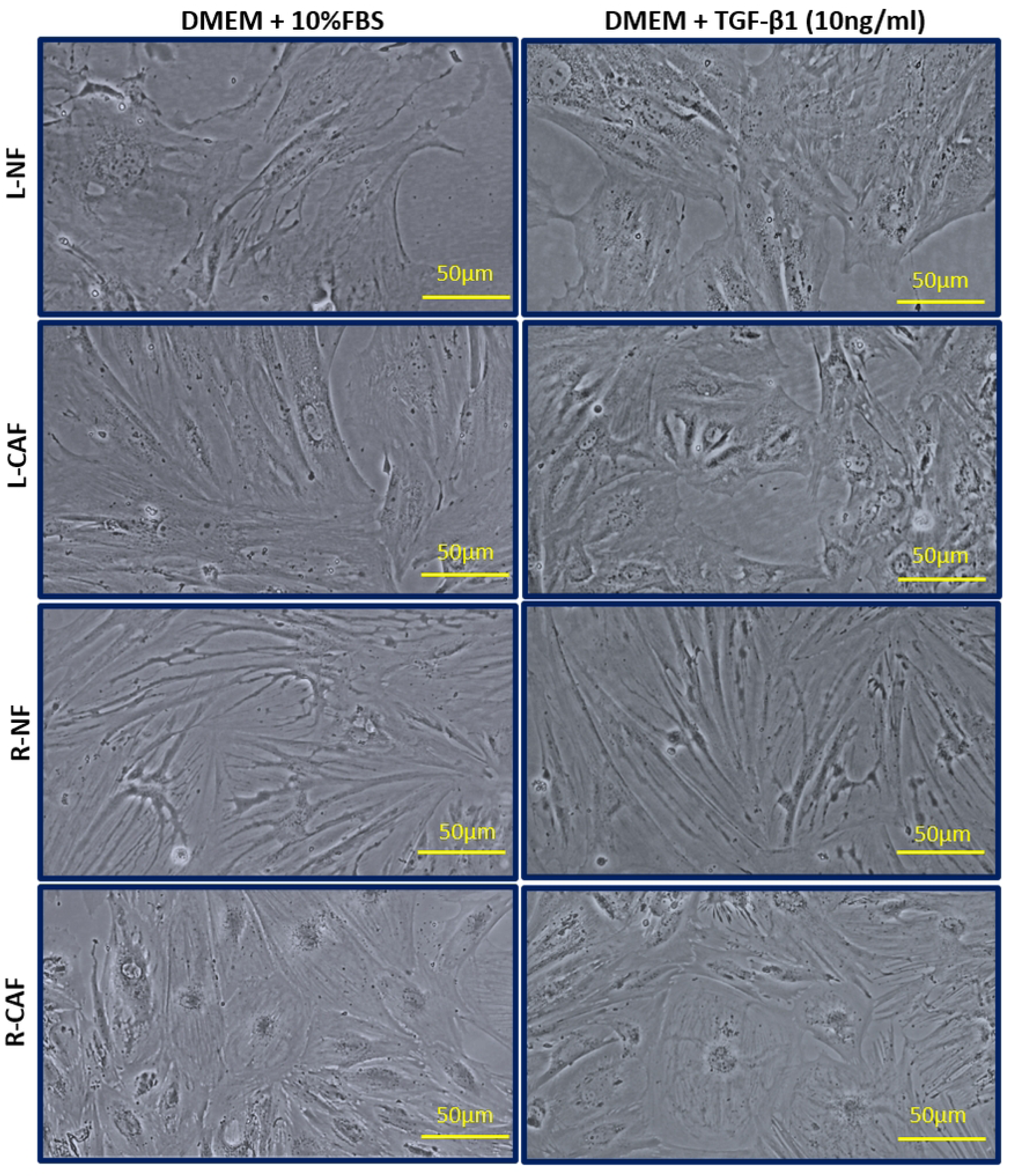
Morphology of fibroblasts observed following 72 hours treatment under DMEM + 10%FBS and DMEM + TGFβ1 (10ng/ml).

## Discussion

The present study reveals distinct profiling of colorectal fibroblasts in the TME of LCRC and RCRC. Transcriptomic profiling identified several top DEGs between primary fibroblasts isolated from CRC tissues and SW620 cells, which are fibroblast-specific and associated with ECM functions [11–16]. *EpCAM* expression in epithelial cancer cells except fibroblasts also confirm the fibroblast phenotype of the established primary cell lines in the present study. Among these DEGs, *RBPMS* was notably upregulated in fibroblasts. Previous studies have reported RBPMS expression in other cancer types [17, 18]. Gyorffy [19] found that RBPMS upregulation along with tissue inhibitor of metalloproteinases 1 (TIMP1), COL4A2, and transgelin (TAGLN) which are synonyms of CAFs, correlates with shorter survival in CRC. Worth noting that most studies focused on RBPMS expressing epithelial cells rather than fibroblasts of the TME.

Our data strongly highlights the specific expression of RBPMS which enables us to distinguish L-CAFs and R-CAFs. RBPMS, reported as a transcriptional co-activator, has known roles in smooth muscle cells differentiation and cardiac function [20, 21]. RBPMS upregulation promotes epithelial-mesenchymal transition (EMT) and tumor desmoplasia activity [18, 22, 23], and is associated with high lymph node metastasis in HDAC2 mutant CRC and gastric cancer cells [23, 24]. Conversely, Rabelo-Fernández et al. [25] reported that reduced RBPMS expression drives tumor proliferation by repressing AP-1 transcriptional activity. This could partially explain the RBPMS downregulation in R-CAF. Loss of RBPMS is also associated with resistance to EGFR inhibitors through enhanced HER2/AKT/mTOR signaling [26]. As RBPMS exists in multiple isoforms (RBPMS/RBPMS1 (30 kDa), RBPMS2 (22.4 kDa), and RBPMS3 (24.2 kDa)) arising from alternative splicing [25], further investigation is needed to determine which RBPMS is expressed differently in the fibroblasts.

Interestingly, the observed RBPMS specific expression correlates with key CAF subtypes that further distinguish L-CAF and R-CAF. These CAF subtypes include myofibroblastic CAFs (myCAFs) and inflammatory CAFs (iCAFs), both of which are involved in fibroblast activation [7]. L-CAF exhibited high levels of α-SMA, myofibroblasts, and ECM related markers, hence defining them as myCAFs whereas, R-CAF showed a lower level of α-SMA (*ACTA2*) and high levels of inflammatory markers, indicative of iCAFs. To our knowledge, the present study is the first to report such CAF subtypes in the context of CRC sidedness. Previously, Öhlund et al. [27] reported a groundbreaking discovery in classifying CAFs from pancreatic cancer into either myCAFs or iCAFs based on their molecular phenotypes. The myCAF populations are in close proximity to tumor cells, express a high level of α-SMA, and are stimulated by TGFβ whereas, iCAFs are located at a considerable distance from tumor cells, with an extremely low level of α-SMA and elevated production of inflammatory cytokines (e.g., IL-6 and leukemia inhibitory factor). Additionally, AOC3 downregulation and LRRC17 upregulation observed in L-CAF are consistent with myCAF phenotype [8, 9].

MyCAF is linked to highly invasive tumors with high lymphatic invasion [28]. This could reflect the underlying desmoplastic characteristic of LCRC tumors as reported in Wang et al. [29] with more tumor invasion-related characteristics in the fibroblasts of LCRC. Consistently, ECM markers (*IGFBP5* and *MMP2)* support tumor invasion and metastasis [30, 31]. In contrast, iCAFs drive tumor proliferation and angiogenesis [32]. The greater proliferation of R-CAF compared to L-CAF supports the aggressive iCAF phenotype, which may underline the more aggressive behavior of RCRC tumors. The chemokine *CXCL2* has been reported to promote proliferation in colon cancer cells [33], supporting the increased SW620 cells proliferation upon incubation with CM of R-CAF. *CCL7* and *CCL14*; *CCL15* are also implicated in tumor migration and associated with poor prognosis [34, 35]. Additionally, *THBS1,* another ECM marker downregulated in R-CAF has been shown to promote pro-angiogenic activities in RCRC [36].

Further analyses revealed that RBPMS expression is TGFβ-dependent, suggesting a potential interplay between these two proteins in shaping CAFs and CRC heterogeneity. Prior to TGFβ1 treatment, transcriptomic profiling identified the higher levels of both *RBPMS* and *TGFβ1* in L-CAF compared to R-CAF. RBPMS upregulation maintains the myCAF hallmark in TGFβ1-treated L-CAF, as supported by AOC3 downregulation, and the upregulation of both LRRC17 and α-SMA. MyCAF transdifferentiation is also observed in TGFβ1-treated L-NF. This suggests that the TGFβ-dependent RBPMS may reinforce the myCAF transdifferentiation while, its loss in R-CAF, may support the aggressive iCAF phenotype. However, this relationship is multifactorial as CAF activation is modulated by cytokines, myofibroblast, and ECM markers [6, 7]. Hence, further studies are needed to clarify the functional role of RBPMS in CAF-regulated phenotype.

Primarily, RBPMS is implicated in TGFβ signaling, a regulatory pathway that functions as both a tumor-suppressor and a tumor-promoter, depending on the cellular context [37, 38]. RBPMS upregulation promoted SMAD2-4 phosphorylation which is correlated with EMT-phenotype [39]. Consistent with TGFβ’s role as a tumor suppressor, RBPMS knockdown reduced SMAD3/4 activity, leading to pro-proliferative and pro-angiogenic behaviors associated with poor prognosis in ovarian cancer [25]. Hence, the TGFβ1 regulation of RBPMS could be site-specific. The CAF-RBPMS/TGFβ1 signaling may operate through distinct feedback mechanisms, with a positive loop in LCRC promoting tumor desmoplasia and a negative loop in RCRC contributing to tumor aggressiveness.

As for AOC3 heterogeneous expression in the TGFβ1-treated fibroblasts, this also suggests the regulation of AOC3 by TGFβ1. Though, little is described on AOC3 relationship with TGFβ1, hence require further investigation.

Our study also revealed the importance of bidirectional CAF-cancer cell crosstalk in CAF-regulated phenotype and carcinogenesis. Previous studies support the significant pro-proliferative effects of CAF secretomes on SW620. Herrera et al. [40] reported increased proliferation of CRC cells (SW480) with Snail1-overexpresing from CM of CAF. You et al. [41] also observed CAF-platelet-derived growth factor (PDGF) mediated proliferation of oral squamous cell carcinoma. However, some fibroblasts subgroups suppress colonic tumor growth [42], supporting the inhibitory effects of NF secretomes on SW620 proliferation.

Although L-CAF and R-CAF may differ morphologically which suggest phenotypic differences, this was inconclusive given that both CM of L-CAF and R-CAF induced SW620 proliferation. This indicates that CAFs drive cancer proliferation regardless of colon sidedness.

Nevertheless, for further validation of RBPMS expression, more CRC cell lines and fibroblasts from similar or varying contexts, as well as other cytokines are needed to better elucidate its specific expression and regulation by TGFβ1 in colorectal carcinogenesis. Gene knockdown experiments targeting those markers (*RBPMS*, *α-SMA*, *AOC3*, and *LRCC17*) to dissect the signaling pathways involved in CAF activation would be a future direction to pursue. Further characterization of the influence of CAFs with high or low expression of RBPMS on bidirectional communication with different subtypes of CRC cells (i.e., differentiated versus undifferentiated) would be vital in dissecting the role of this marker in colorectal carcinogenesis. Moreover, the ability of high- and low-RBPMS-expressing CAFs to promote CRC *in vivo* would be essential to be investigated using animal models. These would provide further insights related to protein-protein interaction and regulation in heterogeneous CRC biology.

Recent work using single-cell RNA sequencing has identified functional heterogeneity in MYH11+ CAF subtypes between LCRC and RCRC [29]. Our study corroborates the CAFs heterogeneity in the context of CRC sidedness, as demonstrated by the novel discovery of RBPMS as a potential CAF marker and the enrichment of distinct CAF subtypes (i.e. myCAFs and iCAFs). With its regulation by TGFβ1 signaling, RBPMS may serve as a target in CAF-based therapy for CRC sidedness. Several efforts in targeted-CAF based therapy have been proposed to improve chemotherapy or immunotherapy efficacy. Qi et al. [43] have devised a fibroblast activation protein, alpha (FAPα)-activated prodrug Z-GP-DAVLBH, which functions to deplete FAP+CAFs, potentially reducing the bevacizumab-resistance in metastatic CRC patients. Another study reported that targeting CAF-derived IL8 improved anti-PD-L1 effects in alleviating tumor immunosuppression [44]. Although CAF-targeted therapy is still under investigation, its impact on CRC suggests a promising avenue for enhancing treatment outcomes and improving patient survival.

## Conclusion

RBPMS was identified as a specific marker expressed among fibroblasts from different sidedness of the human colon, exclusively in CAFs isolated from LCRC and RCRC tissues. Moreover, RBPMS regulation by TGFβ1 was demonstrated, along with the identification of myCAFs in LCRC and iCAFs subtypes in RCRC. Therefore, RBPMS may serve as a target in CAF-based therapy in the future and offers more insights into the mechanisms differentiating LCRC and RCRC.

## Acknowledgements

The authors would like to acknowledge Dr Nazihah Mohd Yunus and Nur Sabrina Abdul Rashid for their support in the present study.

